# A novel single-cell RNA-sequencing platform and its applicability connecting genotype to phenotype in ageing-disease

**DOI:** 10.1101/2021.10.25.465702

**Authors:** Orr Shomroni, Maren Sitte, Julia Schmidt, Sabnam Parbin, Fabian Ludewig, Gökhan Yigit, Laura Cecilia Zelarayan, Katrin Streckfuss-Bömeke, Bernd Wollnik, Gabriela Salinas

## Abstract

Advances in single cell multi-omics analysis such as the ability of a genotype to show phenotypic diversity lead to a comprehensive understanding of cellular events in human diseases. The cardinal feature to access this information is the technology used for single-cell isolation, barcoding, and sequencing. While numerous platforms have been reported for single-cell RNA-sequencing, technical enhancements are needed in terms of highly purified single cell selection, cell documentation or limitations in cell sizing and chemistry. In this study, we present a novel high-throughput platform using a full-length RNA-sequencing approach that offers substantial technical improvements of all parameters. We applied this platform to dermal fibroblasts derived from six patients with different segmental progeria syndromes and defined phenotype-associated pathway signatures and variant-associated expression modifiers. These results validate the applicability of our platform to highlight genotype–expression relationships for molecular phenotyping in ageing disorders.

## Introduction

In the field of genomic medicine, contemporary single-cell sequencing methods are focusing on the characterization of individual cells. Recent advances in integrated robotic systems and molecular barcoding have made the transcriptional profiling of thousands of individual cells cost-effective^1,2^.

Available integrated systems for single-cell applications differ in their sensitivity, specificity and throughput. They also differ in other critical experimental parameters such as the detection of doublet and debris, limitations in capturing rate due to cell sizing or cell morphology and documentation or visualization of the captured single cells.

The most critical step for obtaining transcriptome and genome information from individual cells is the single cell isolation, and thus it is necessary to distinguish between methods with low- or high-throughput for the collection of single cells. Low-throughput approaches such as limiting dilution, micromanipulation and laser capture microdissection (LCM) are time-consuming and exhibit limitations in the capture of rare cells^2,3^. Flow-activated cell sorting (FACS) is a commonly used strategy for isolating highly purified single cells. The potential limitations of these techniques include the requirement of large starting volumes (difficulty in isolating cells from low-input numbers <10,000) and the need for monoclonal antibodies to target proteins of interest leaving novel cell populations unexplored^2–4^.

Recently, integrated systems for both single cell collection and downstream experiments, particularly those related to single-cell NGS-applications have become commercially available. In particular, the microdroplet-based microfluidics technology such as those used by Drop-Seq and 10x Genomics Chromium platform, uses a gel bead coated with oligonucleotides^5,6^. These platforms present some important limitations including the use of cell suspension dilution for cell capturing, cell sizing (< 35 µm), as well as the inability to visually inspect the collected cells and perform full-length cDNA sequencing. Previously, several studies explored the utility of the ICELL8® platform for single-cell sequencing in the cardiac field in which the size of cardiomyocytes limits the use of drop-seq and other platforms^7,8^. The ICELL8® technology allows the sequencing of nuclei and cells with a large range of sizes (3 µm to -500 µm in diameter) and its chemistry is compatible for both living as well as fixed cells. Depending on the biological needs, different ICELL8® Chips and chemistries are available for RNA-sequencing (RNA-Seq) and downstream applications.

Several groups performed comparisons between different single-cell RNA-sequencing (scRNA-seq) platforms and chemistry^9–12^. However, these comparisons utilized custom-build drop-seq platforms and/or reaction chemistry, which can potentially be a hurdle for investigators new to the scRNA-seq field^13–15^.

In this paper, a combination of two platforms, the CellenONE® X1 (CellenONE) and the ICELL8® cx Single-Cell System (ICELL8), was implemented and validated as a novel method for scRNA-sequencing. The CellenONE allows the selection of highly purified single cells based on cell sizing, cell morphology or by using one or more fluorescence markers. Moreover, we improved the cell capturing efficiency to nearly 80% by collecting cells from the CellenONE directly into a Takara ICELL8® 5,184 nano-well chip. Here we report a single-cell full-length RNA-seq approach that uses the SMART-Seq technology in the single-cell platforms, demonstrating the molecular characterization of mutational and transcriptional heterogeneity in dermal fibroblasts derived from six patients with different segmental progeria syndromes. In addition to the causative mutations previously identified in these patients^16–19^, we performed regulon analysis to identify interactions of genetic alterations that can be held responsible for the phenotypic heterogeneity.

## Results

### Method overview of integrated platforms for scRNA-sequencing

Figure 1 compares the most used state-of-the-art integrated platforms for single-cell collection. All platforms have an integrated system that includes cells dispensing, conversion of RNA into cDNA (RT) and library preparation for sequencing. The microfluidic droplet (Figure 1a) is able to process thousands of individual cells in a fully automated manner. However, it is lacking the capability of evaluating the quality of single cells based on imaging features, discarding damaged cells that potentially generate artificial cell clusters. Moreover, the chemistry used is based on 3’ or 5’ fragment, which is not optimal for the investigation of mutational heterogeneity in single cells.

**Figure 1:**
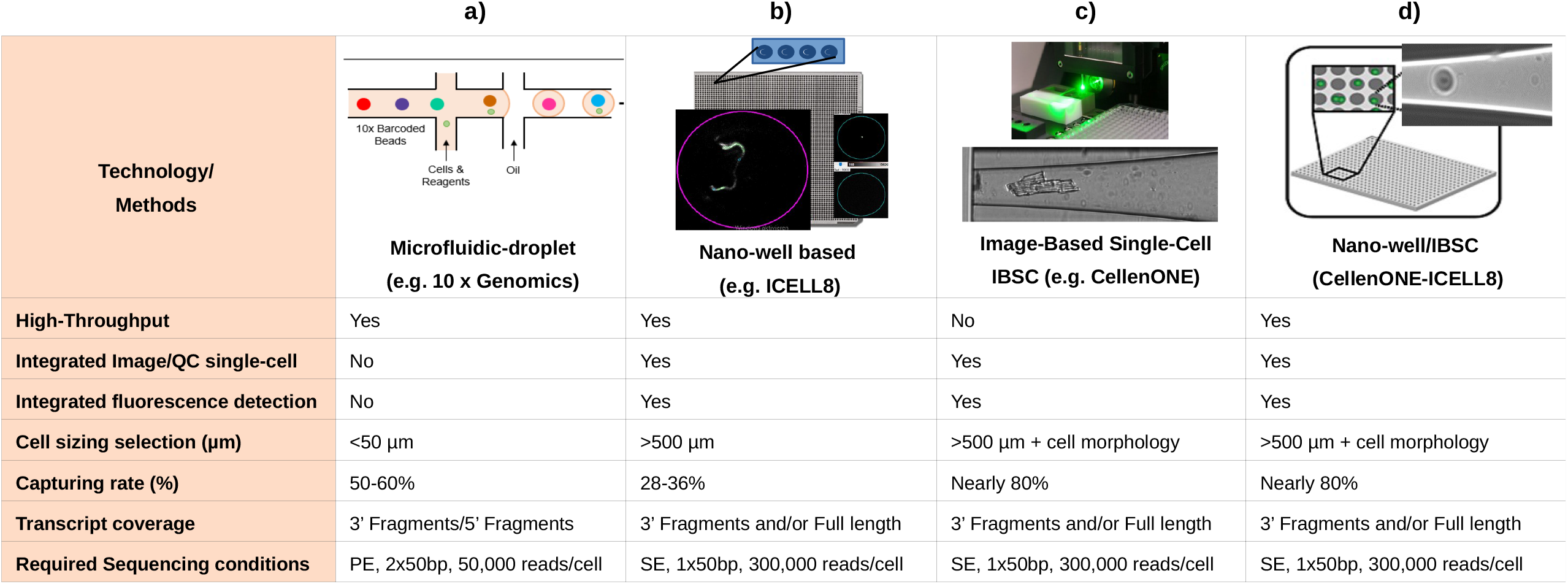
Technical information of state-of-the-art integrated platforms for single-cell RNA-Sequencing. Methods used in each of the platforms for single-cell isolation, barcoding, chemistry and sequencing. (a) the 10 x Genomics Chromium (b) the ICELL8 cx Single-Cell system; ICELL8 (c) the CellenONE® X1 system, CellenONE and (d) advantages of the combined CellenONE® X1 and iCELL8 cx Single-Cell systems, CellenONE-ICELL8.

The ICELL8 (Figure 1b) can select living cells, while discarding dead cells and doublets by imaging. The instrument exhibits a low cell-capturing rate that varies between 24-36% from 5,184 cells. Depending on the cell system used, a range of 800 to 1400 cells can be analyzed in one experiment (one chip with 5,184 nano-wells). The advantages of the ICELL8 include flexible chemistry for transcript coverage (3’ end fragment or full-length approaches), flexible sequencing mode selection (choosing between single-end SE or paired-end PE mode) and dispensing of cells with abnormal morphology e.g., Schwann cells that exhibit axonic regrowing forms and sizing of 200-500 µm (Figure 1b and supplementary Figure 1).

The CellenONE (Figure 1c) offers an excellent imaging system for cell collection. The technology, which is based on Image Based Single Cell Isolation (IBSCIT™), offers high-definition optics for cell visualization and can isolate cells over a wide size range. For example, the dispensing of cardiomyocytes with a cylindrical shape and dimensions of 35-120 µm can be performed using this system^20,21^. One disadvantage this instrument presents is its inability to simultaneously process thousands of cells for sequencing in a fully automated manner.

In order to improve the capturing rate of the ICELL8 and to simultaneously process more than 3500 cells to libraries for sequencing, we combined both instruments (Figure 1b and 1c) by collecting the cells from the CellenONE directly into a Takara ICELL8® 5,184 nano-well chip (Figure 1d).

The system allows the isolation of living or fixed cells, and the performance of an analysis on nuclei or whole cells without limitations regarding cell sizing or cell morphology. Moreover, the exclusion of cell doublets and debris can be done by defining the parameters during cell collection or by cell inspection after dispensing. Each collected cell is reported and documented allowing an additional check for cell quality before sequencing.

### Validation of scRNA-seq platforms using dermal fibroblasts derived from patients with different segmental progeria disorders

In the frame of the clinical and molecular work in our recently established Center for Progeroid Disorders, we had previously used NGS-based approaches to determine the molecular diagnosis of progeria patients in early stages of a disease^16–19^. Although the identification of primary causative genes and pathogenic variants is essential for understanding the patient-specific pathogenesis, systematic biological screens are necessary to elucidate variant and transcriptional heterogeneity. In order to shed light on additive genetic and transcriptional drivers of premature ageing processes, we used two different scRNA-seq platforms: ICELL8 alone and a combination of ICELL8 and CellenONE. The two platforms were compared and validated for their diagnostic applicability using dermal fibroblasts from a unique cohort from six progeria patients (GOE1360, GOE1309, GOE800, GOE615, GOE486 and GOE247) and two age-matched, healthy control individuals (GOE1303 and GOE1305). Initially, single cells from the eight dermal fibroblasts samples were dispensed by the CellenONE to a single a Takara ICELL8® 5,184 nano-well chip. The dispensing parameters used for human dermal fibroblasts cells were based on sizing (20-60 µm in diameter, elongation and circularity of 1-1.5 µm, see Figure 2a). Following first quality control based on single cell collection, we discarded 123 (doublets) of 5,184 cells. After sequencing and QC filtering, a total of 3,135 cells were processed for data analysis (Figure 2a, Table 1). For the ICELL8 single-cell platform, cells from the same samples were dispensed in three ICELL8® 5,184 nano-well chips obtaining 3,129 cells after data merging and sequencing QC filtering (Figure 2b, Table 1). It is important to note, that the batch effect observed by merging data from three ICELL8® 5,184 nano-well chips dispensing the same samples was very low (supplementary Figure 2), suggesting that a more sensitive platform with deeper sequencing depth can mitigate the variation to a large degree of cells.

**Table 1:** Number of cells dispensed and selected for sequencing after QC-filtering data set (CogentDS).

| **Samples CellenONE-ICELL8** | **GOE1303** | **GOE1305** | **GOE1309** | **GOE1360** | **GOE247** | **GOE486** | **GOE615** |
| --- | --- | --- | --- | --- | --- | --- | --- |
| No. of cells dispensed | 576 | 576 | 576 | 576 | 576 | 576 | 576 |
| No. of cells Post-QC | 387 | 295 | 298 | 537 | 479 | 195 | 218 |
| Total cells Post-QC | **3135** |  |  |  |  |  |  |
| **Samples ICELL8** | **GOE1303** | **GOE1305** | **GOE1309** | **GOE1360** | **GOE247** | **GOE486** | **GOE615** |
| No. of cells dispensed | 436 | 364 | 485 | 356 | 446 | 477 | 350 |
| No. of cells Post-QC | 423 | 323 | 411 | 280 | 386 | 445 | 317 |
| Total cells Post-QC | **3129** |  |  |  |  |  |  |

**Figure 2:**
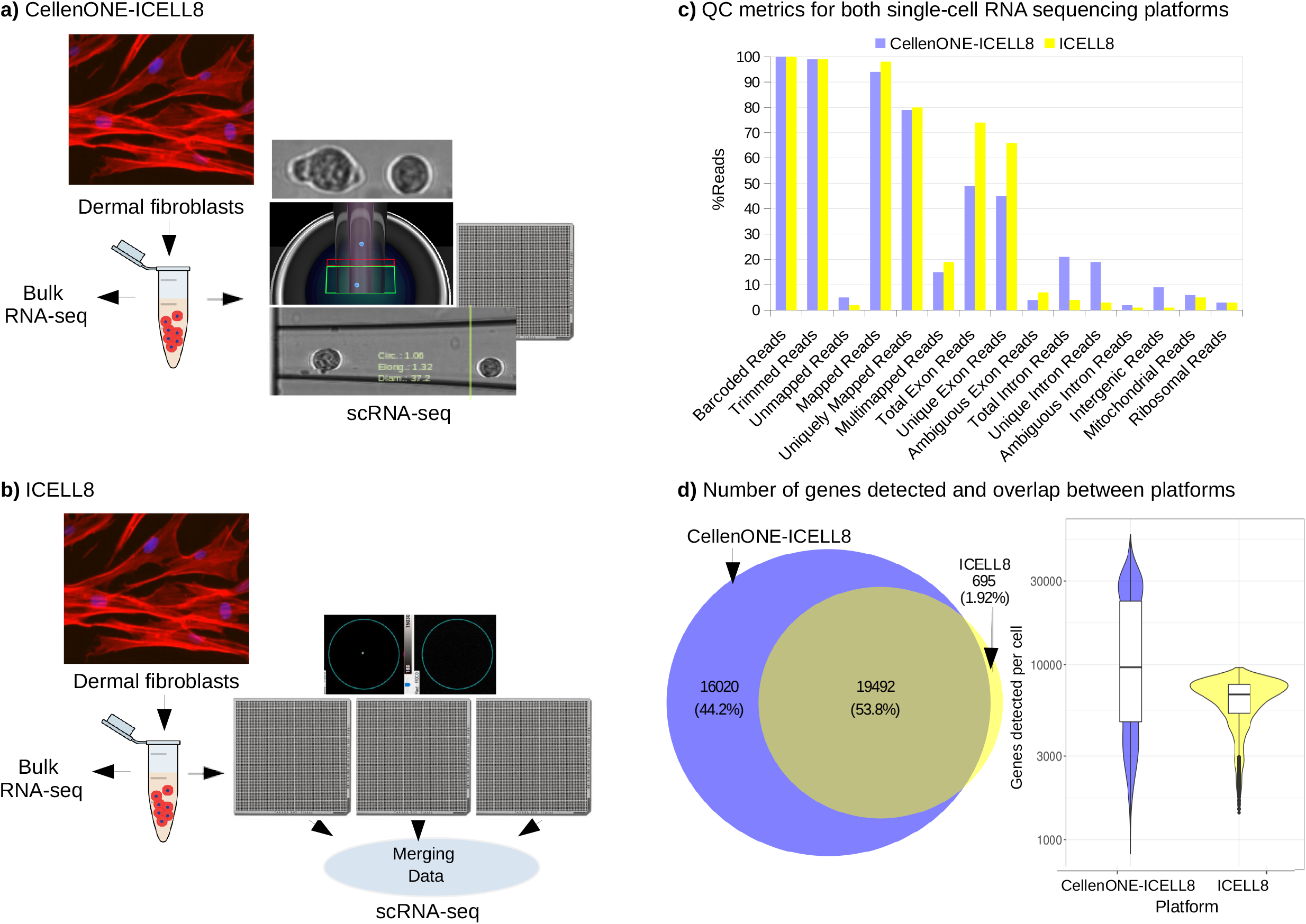
Study design for the scRNA-Seq platforms validation. (a) Experimental design for the CellenONE-ICELL8 Single-Cell System combination. Only one ICELL8® nano-well chip was used for all eight dermal fibroblasts patient samples. (b) Experimental design for the ICELL8 Single-Cell System. A total of three (3) ICELL8® nano-well chips were used for all eight dermal fibroblasts patient samples. (c) Bar plot displaying proportions of reads left in both single-cell platforms after each step of the analysis. (d) Venn diagram displaying the number of genes in QC-filtered data from both approaches and the violin plot shows the number of QC-filtered genes expressed in individual cells in both approaches.

The datasets generated from both platforms were comparable in terms of numbers of total reads (1.01G to 1.67 G) and reads per barcode (272K to 309K). Interestingly, when comparing both approaches, the dataset generated in the combined platform showed a higher percentage of intronic and intergenic reads than the ICELL8 (Figure 2c, Table 2a). Furthermore, across all samples investigated, the portion of non-coding RNAs detected by the combined platform, especially the detection of long non-coding RNAs (lncRNAs), was significantly higher than those detected in the ICELL8 platform. The biotype and Table 2b show an example for sample GOE1309 with 29% of the dataset; when using the combined platform compared to 16% on the ICELL8 platform. This suggests that while the novel approach results in sufficient amount of exonic reads, it may also provide additional insight into the utility of intergenic sequences and introns (for instance intronic retention) on an individual cell level.

**Table 2a:** Reads parameters of the two scRNA-sequencing platforms CellenONE-ICELL8 and ICELL8 (CogentDS).

| QC-Parameters | CellenONE-ICELL8 | ICELL8 |
| --- | --- | --- |
| Samples | 8 | 8 |
| No. of cells | 5184 | 3,616 cells |
| No. of cells Post-QC | 3135 | 3,129 cells |
| Total Reads | 1.67 G | 1.01 G |
| Barcoded Reads | 1.61G | 985 M |
| Fraction Barcoded Reads | 0.96 | 0.98 |
| Barcodes Identified | 5,184 cells | 3,616 cells |
| Reads per Barcode | 309.84 K | 272.54 K |
| Barcoded Reads | 1606229682 (100%) | 985522031 (100%) |
| Trimmed Reads | 1601535800 (99.71%) | 982927397 (99.74%) |
| Unmapped Reads | 86672148 (5.4%) | 16885303 (1.71%) |
| Mapped Reads | 1514863652 (94.31%) | 966042094 (98.02%) |
| Uniquely Mapped Reads | 1270205342 (79.08%) | 778278333 (78.97%) |
| Multimapped Reads | 244658310 (15.23%) | 187763761 (19.05%) |
| Total Exon Reads | 788938677 (49.12%) | 727757440 (73.84%) |
| Unique Exon Reads | 714643829 (44.49%) | 654119292 (66.37%) |
| Ambiguous Exon Reads | 74294848 (4.63%) | 73638148 (7.47%) |
| Total Intron Reads | 337929018 (21.04%) | 37383326 (3.79%) |
| Unique Intron Reads | 300416455 (18.7%) | 32630018 (3.31%) |
| Ambiguous Intron Reads | 37512563 (2.34%) | 4753308 (0.48%) |
| Intergenic Reads | 143337647 (8.92%) | 13137567 (1.33%) |
| Mitochondrial Reads | 93167354 (5.8%) | 57107478 (5.79%) |
| Ribosomal Reads | 30369153 (1.89%) | 29861198 (3.03%) |

**Table 2b:** RNA-Biotype percentages identified in sample GOE1309 using the CellenONE-ICELL8 and ICELL8 technologies.

| RNA Biotype | CellenONE-ICELL8 (%) | ICELL8 (%) |
| --- | --- | --- |
| Coding genes | 49 | 72 |
| lncRNA | 29 | 16 |
| Processed_pseudogene | 12 | 8 |
| Unprocessed_pseudogene | 3 | - |
| TEC | 3 | - |
| Transcribed_pseudogene | 2 | 4 |
| Misc_RNA | 2 | - |

Comparing the QC-filtered transcribed genes from both methods has shown that 52% of all genes overlapped between the approaches, 1.5% were ICELL8-specific and 46% were specific in the combined approach (Figure 2d). This result demonstrates that the novel platform can reproduce sequencing data generated by ICELL8, and has the advantage of providing additional genetic information with increased coverage of more genes.

### Correlations of gene expressions between single-cell platforms and bulk RNA-seq

To evaluate similarities in gene expressions between the single-cell approaches, normalized expressions from both single-cell datasets were correlated for the top 100 markers for each sample (Figure 3a). The correlations appeared high, ranging from 0.6 to 0.85 between the methods, providing additional evidence of the similarity in gene expression, even across different samples.

**Figure 3:**
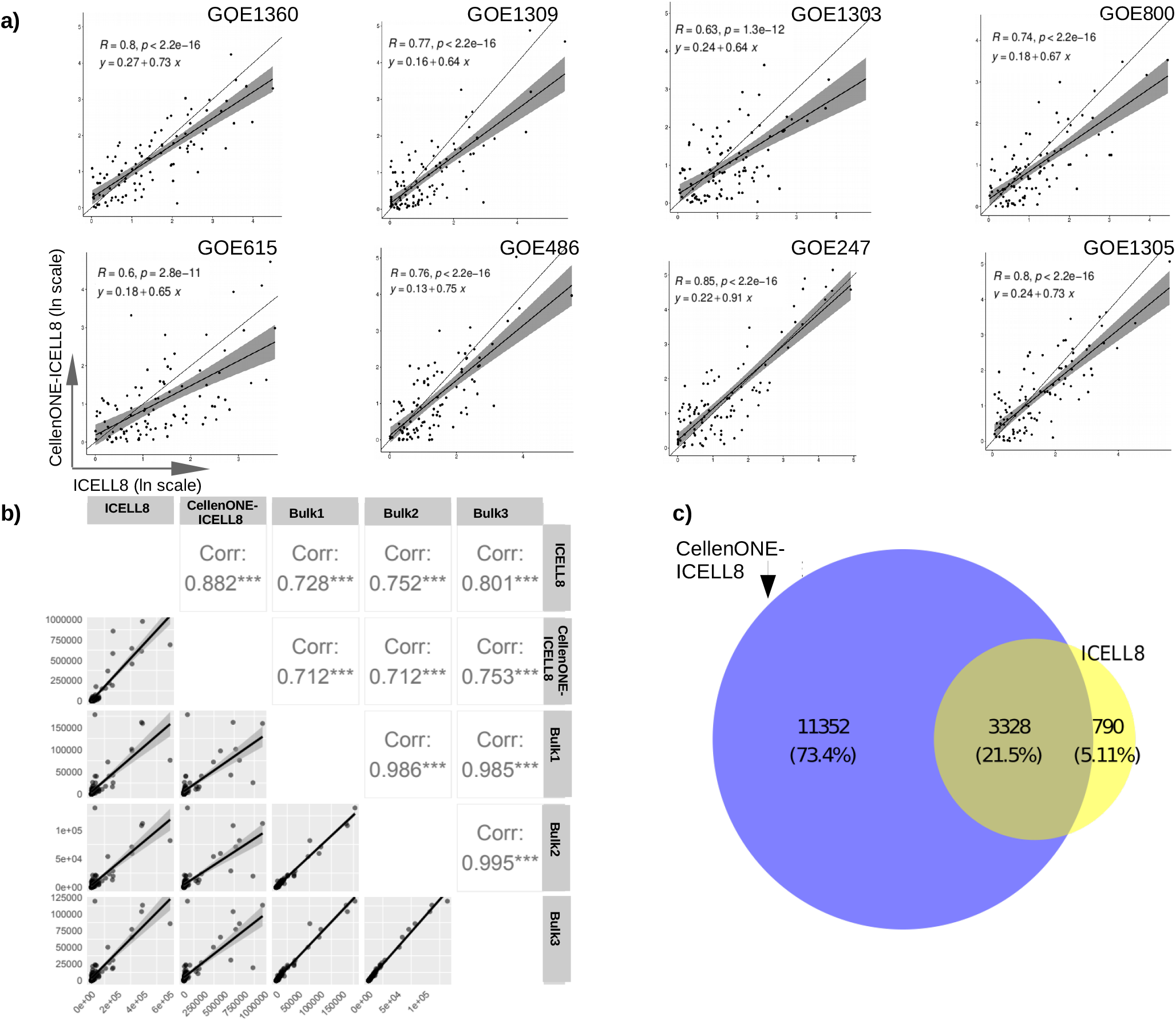
Correlations of scRNA-Seq platforms and bulk RNA-seq. (a) Normalized expressions of top 100 markers in each sample from both single-cell platforms: ICELL8 and CellenONE-ICELL8, including a regression line with 95% confidence interval (in gray) and the overall correlation coefficient R. (b) Normalized expression and correlation of the top 100 markers in sample GOE247 between both single-cell and bulk RNA datasets, with the correlation coefficients indicated in the upper triangle (c) Venn diagram of all significant markers detected for sample GOE247 in both platforms (ICELL8 and CellenONE-ICELL8).

Moreover, by looking at the markers detected for individual samples (described in Figure 5), we observed a significant overlap between the platforms. The CellenONE-ICELL8 approach shows many additional unique markers that the ICELL8 alone does not detect (Figure 3c for sample GOE247, Supplementary Figure 4 for other samples). This correlates with the results in Figure 2d, indicating high sensitivity on gene detection by the performance of the new platform and a significant overlap between both methods. We also performed bulk RNA-sequencing using the same eight fibroblasts’ samples selected for scRNA-seq, which was used as a reference on the sensitivity on gene detection of the scRNA assay. Normalized pseudo-counts from the single-cell approaches were correlated with the normalized counts for the bulk RNA-seq data using the top 100 markers mentioned above. For example, the correlation for sample GOE247 ranged from 0.71 to 0.99 (Figure 3b; all other samples are shown in Supplementary Figure 3). Overall, the analysis between scRNA-seq platforms and the bulk RNA-seq data showed a high correlation and large overlap in markers, suggesting a strong agreement on the genomic profiles of cells/samples between these approaches.

**Figure 4:**
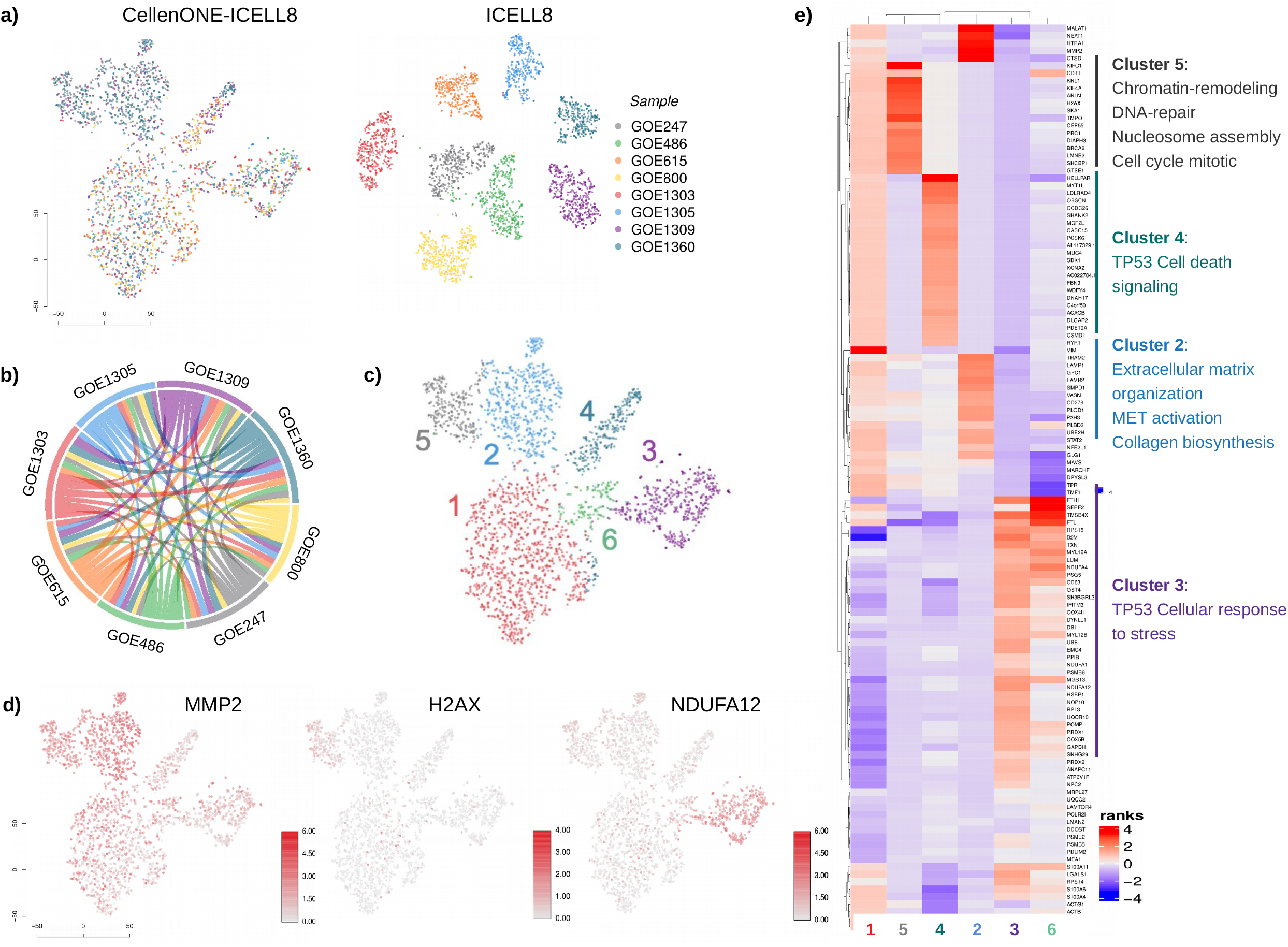
Analysis of transcriptional heterogeneity of dermal fibroblast derived from patients using the novel platform. (a) t-SNE of the ICELL8 and CellenONE-ICELL8 experiments. (b) Circo plot showing connection of top 100 marker genes of the samples in both ICELL8 and CellenONE-ICELL8 platforms. (c) Unsupervised t-SNE of the CellenONE-ICELL8 approach. (d) t-SNE showing most prominent markers for different clusters. (e) Heatmap of rank scores of the top 30 markers in each unsupervised cluster, including enriched pathways.

**Figure 5:**
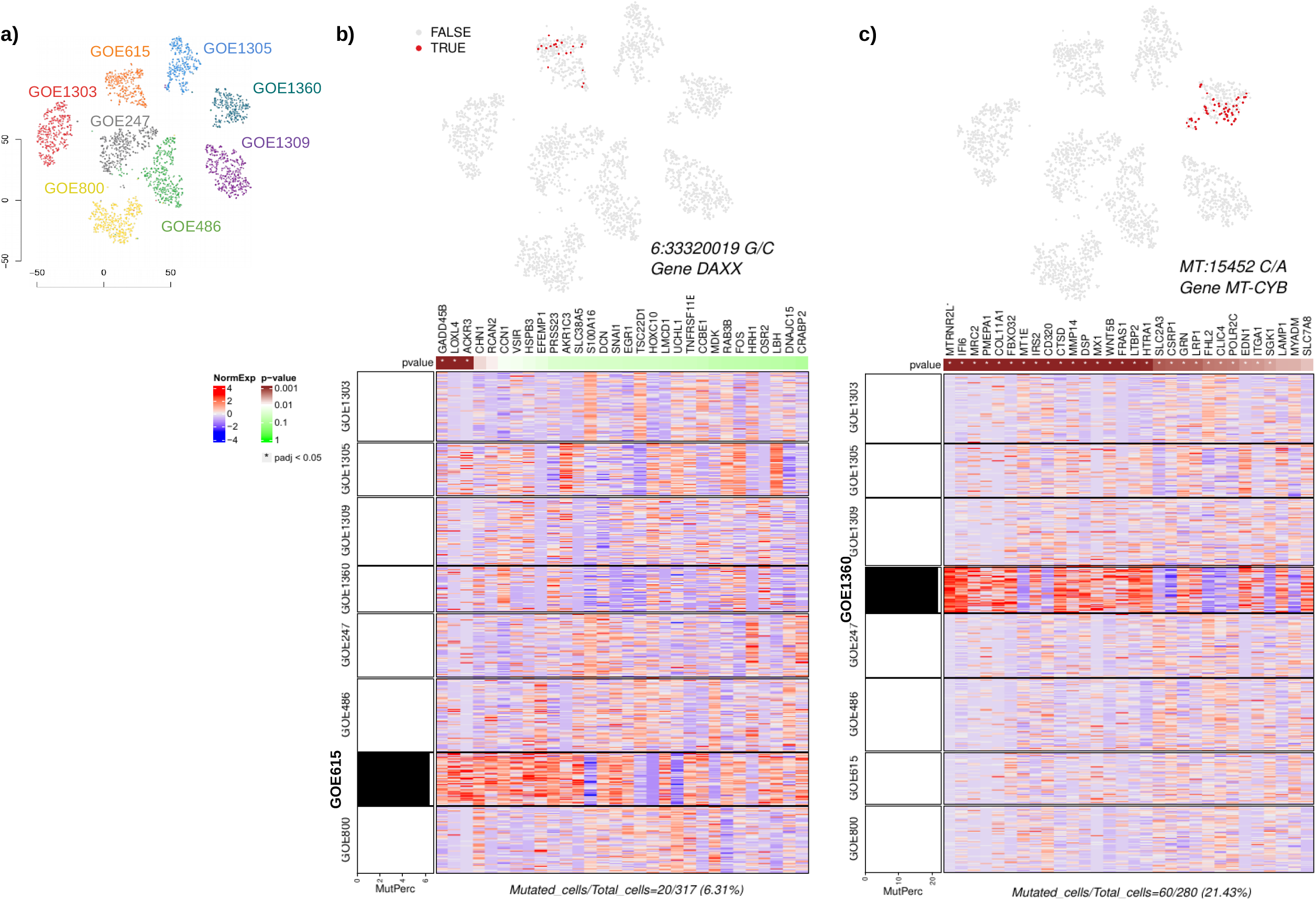
Analysis of mutational heterogeneity in dermal fibroblast derived patients. (a) supervised clustering showing expression of single cells. (b) t-SNE projection showing non-uniformly distributed DAXX^A486G^ variant-expressing cells in patient GOE615 in red and the regulon for variant-dependent genes. The heatmap displays the scaled, normalized gene expressions of the top 30 markers with highest adjusted p-values from the regression of the gene expressions to the mutation rate (p-values shown in top bar), and a bar graph on the leftt shows mutant cell fraction in each cluster labelled to the left of the heatmap. (c) t-SNE projection showing non-uniformly distributed MT-CYB^L236I^ variant-expressing cells in GOE1360 in red and the regulon for variant-dependent genes, with a bar graph showing mutant cell fraction in each cluster labelled to the left of the heatmap.

### Determination of the transcriptional heterogeneity in accelerated ageing phenotypes

In order to validate the scRNA seq platform and to shed light on the transcriptional signatures we used the very unique patient cohort, all representing premature ageing phenotypes. All of these samples has been described previously on cellular hallmarks of ageing driving physiological and pathological ageing processes, e.g. such as genomic instability, telomere attrition, epigenetic alterations, loss of proteostasis, mitochondrial dysfunction, deregulated nutrient sensing, stem cell exhaustion, cellular senescence and altered intercellular communication^22,23^.

Previous data from gene panel and whole exome sequencing approaches identified causative mutations in these patients: GOE615 (*PYCR1*, c.797+2_797+5del [homozygous]), GOE1360 (*LMNA*, c.1821G>A^16^), GOE800 (*YRDC*, c.662T>C [homozygous], p.Ile221Thr^19^), GOE486 (*AHCTF1*, c.6329C>G [de novo], p.Ser2110Cys (Unpublished data), p.Lys215_Asp319del^17^), GOE247 (*SLC25A24*, c.593G>A [de novo], p.R198H^18^), GOE1309 (*BLM*, c.1544dup, p.Asn515Lys^*^2 and c.2254C>T, p.Gln752^*^ [compound heterozygous] (Unpublished data).

For dimensionality reduction and data visualisation we performed the t-SNE algorithm (as implemented in CogentDS). To evaluate transcriptional events regarding samples in both scRNA-seq methods, we analysed cell groups based on their sample association (Figure 4a) and performed, in parallel, unsupervised graph-based clustering of cells based on their gene expressions (Figure 4c).

The unsupervised clustering displayed six clusters with the most significant differences in gene expression signatures between each cluster (Figure 4c and 4d). The gene profiles evaluated for the clusters were associated to specific markers of dermal fibroblasts patient samples and are related to pathogenesis of different ageing-disorders (Figure 4d and 4e). It is important to note, that several markers appear to be expressed in all samples, indicating transcriptional events that are commonly expressed within all samples. This is reflected in Figure 4b showing the half of the top 100 markers expressed across investigated samples including the controls.

We next compared cluster 1, which predominantly represents the control samples GOE1305 and GOE1303 to all other clusters related to ageing-disorders. Starting with cluster 2, markers related to extracellular matrix organization, MET activation, and collagen biosynthesis (MMP2, P3H3, PLOD1, and LAMB2) were found to be upregulated in comparison to Cluster 1. Cluster 3 shows the over-expression of markers for cellular response to stress and TP53 regulating metabolic genes (NDUFA4, UBB, RPS18, HSBP1, COX4l1, ATP6V1F, TXN, PRDX1, PRDX2, PSMB6, RPL3, COX5B, and ANAPC11) when comparing to controls. Cluster 4 exhibits over-expression of markers associated with cell death signalling via NRAGE, NRIF, NADE, cell death receptor signalling pathway (MCF2L, OBSCN) and activation of metabolic gene expression (ACACB). Notably, cluster 5 shared regulation of several genes with cluster 2, which represented mainly patient samples GOE1360 and GOE247. For cluster 2 and 5 we found upregulation of genes related to chromatin-remodelling, DNA-repair, nucleosome assembly, cell cycle mitotic/checkpoints, deposition of new CENPA-containing nucleosomes, and signalling by Rho GTPases e.g., CDT1, KNL1, TMPO, GTSE1, H2HX, BRACA2, and SKA1. Based on those differing markers found for the different clusters, it is possible to display genes that, while they may not be markers for all cells within a sample, they do show particular patterns that can be linked with different progeria phenotypes.

### Variant-associated expression signatures in accelerated ageing phenotypes

We next investigated the extent to which additional variant heterogeneity in putative driver genes could be associated with transcriptional heterogeneity for ageing processes in each case. It is important to note that this analysis was done in a hypothesis-free way, not taking into account the underlying primary genetic cause and mutation of the ageing phenotype, and thereby, being able to search for additive and overlapping variant signatures and expression profiles associated with general aspects of premature ageing.

In order to evaluate how genetic regulation drives specific ageing phenotypes, we searched for patient-specific variants or non-uniformly distributed variants expressed in each of the six patients with different segmental progeria syndromes.

We first combined all variant calling files (VCFs) from both single-cell and bulk RNA-seq data, initially detecting variants in each of the scRNA-Seq samples. Most of the variants were found uniformly distributed, suggesting common effects across all samples related to ageing disease (SNPs and INDELS). Because the detection of SNPs on the scRNA-seq produces many false positives and many of the variants exhibited relatively low depth (1 or 2 reads with a variation), we selected additional criteria to label “mutant cells” considering filtering based on variant- and total depth.

To optimize depth cutoffs, the quartile distributions of the variant and total depths were calculated for each sample. Variants were filtered for variant depths >4 (i.e., number of reads containing the variant base) and total depths >7 (i.e., number of all reads at variant position) in the scRNA-seq data. All variants reported in this study have later been corroborated by the bulk RNA-seq data while filtering variants in the bulk data for variant depths >6 and total depths >10. This resulted in 756 variants for GOE1309, 94 variants for GOE1360, 254 variants for GOE247, 345 variants for GOE486, 273 variants for GOE615 and 234 variants for GOE800 (Supplementary Table 1).

We next evaluated markers whose regulation on expression are influenced by one or more variants by applying a regression-based method^24^. Table 3 described the variants selected for each of the six progeria samples, exhibiting a non-uniformly distributed mutation pattern. All six variants are non-synonymous mutations showing changes in the protein sequence, five of them were found to be reported on the dbSNP and four of them were found to be located on mitochondrial genes.

**Table 3:** Overview of mutations detected in the scRNA-seq and bulk RNA-seq

| Sample | Gene | dbSNP | Chr | Position | Genotype | Codon change | Protein change | Variant name |
| --- | --- | --- | --- | --- | --- | --- | --- | --- |
| GOE615 | DAXX | rs146304558 | 6 | 33320019 | 1/1 | c.1457C>G | p.Ala486Gly | DAXX<sup>A486G</sup> |
| GOE247 | MRPL34 | rs201529220 | 19 | 17306321 | 0/1 | c.221C>T | p.Ala74Val | MRPL34<sup>A74V</sup> |
| GOE800 | MT-CO1 | rs202216551 | M | 6267 | 1/1 | m.6267G>A | p.Ala122Thr | MT-CO1<sup>A122T</sup> |
| GOE1309 | MT-ND4 | rs2853494 | M | 11641 | 1/1 | m.11641A>G | p.Ile294Met | MT-ND4<sup>I294M</sup> |
| GOE486 | MT-ND5 | NA | M | 12557 | 1/1 | m.12557C>T | p.Thr74Ile | MT-ND5<sup>T74I</sup> |
| GOE1360 | MT-CYB | rs193302994 | M | 15452 | 1/1 | m.15452C>A | p.Leu236Ile | MT-CYB<sup>L236I</sup> |

To this end, we highlighted mutant cells on the t-SNE projection of each of the six ageing-samples and evaluated markers related to hallmark mechanisms associated to each phenotype (Figure 5a, 5b, 5c and Supplementary Figure 5). Figure 6b displays variant DAXX^A486G^ (dbSNP rs146304558), which has been identified specifically in sample GOE615. The regulon shows the top 30 markers associated to this specific mutation, where three of them displayed significant regression to the variation with FDR <0.05 (GADD45B, LOXL4 and ACKR3). Among GADD45B related pathways are p53 signaling^25,26^ and DNA damage ATM/ATR regulation of G1/S checkpoint^27^. This protein also associates with centromeres in G2 phase and in the cytoplasm the encoded protein may function to regulate apoptosis^28^.

**Figure 6:**
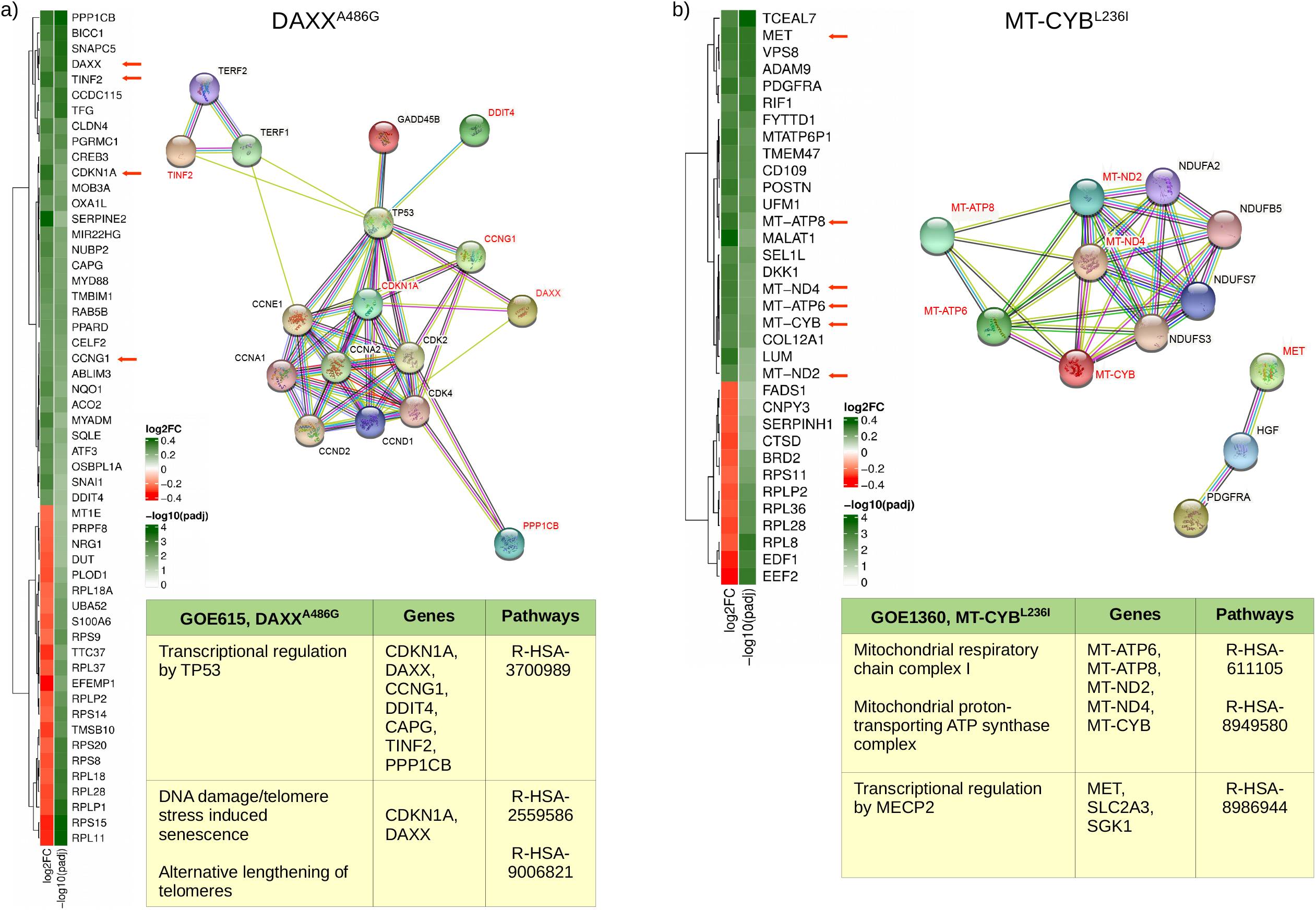
Variant-associated expression signatures in GOE615 and GOE1360. (a) Heatmaps showing log_2_FC and- log_10_ of adjusted p-value for genes differentially expressed in mutant vs non-mutant cells for DAXX^A486G^ variant in sample GOE615 related to protein-protein interaction showing phenotype (StringDB) and pathways (Reactome) for this specific variant. (b) Heatmaps showing log_2_FC and- log_10_ of adjusted p-value for genes differentially expressed in mutant vs non-mutant cells for MT-CYB^L236I^ variant in sample GOE1360 related to protein-protein interaction showing phenotype (StringDB) and pathways (Reactome) for this specific variant.

Notably, four of the sample-specific variants found for GOE800, GOE1360, GOE486 and GOE1309 were located on mitochondrial genes exhibiting high-density mutation in each of the samples (Figure 5c and Supplementary Figure 5). For sample GOE800, variation MT-CO1^A122T^ (dbSNP rs202216551) showed high expression, with six markers showing significant regression to the variation (FDR <0.05). Most of them are involved in hemostasis and formation of fibrin Clot (CD36, SERPINE2, F2R and F2RL2). Specifically, the overexpression of DAB1 was related to Spinocerebellar ataxia 37, epilepsy and involved in the lipoprotein metabolism^29^(Supplementary Figure 5).

Variation MT-CYB^L236I^ (dbSNP rs193302994), found specifically in sample GOE1360, displayed 27 markers as significantly regulated (FDR<0.05) in the heatmap of Figure 5c. Many of those play a role in degradation of the extracellular matrix and collagen degradation (HTRA, LTBP2, COL11A1, ITGA1, MMP14, CTSD, COL11A1), transcriptional regulation by MECP2 (SLC2A3 and SGK1) and neutrophil degranulation (SLC2A3, LAMP1, DSP, GRN, CTSD). In particular, transcriptional regulation by MECP2 has been described in Rett syndrome as rare disease, but still one of the most abundant causes for intellectual disability in females^30^. The type and severity of symptoms are individually highly different. The most important action of MECP2 is regulating epigenetic imprinting and chromatin condensation^30,31^.

In order to understand how different biological pathways explain genetic regulation in association to phenotypes, we used the scRNA-seq data and performed differential expression analysis between cells exhibiting particular variations (mutant cells) and those who do not (non-mutant cells). We specifically performed this analysis on GOE1360 and GOE615 cells containing the MT-CYB^L236I^ and DAXX^A486G^, respectively (Figure 6a, 6b). While in itself a useful method to inspect the effect of variations in cells on gene expressions, we expect its results to coincide with those of the regulon analysis as displayed in Figures 5b and 5c.

For sample GOE615, the differential expression analysis performed between mutant and non-mutant cells for the variant DAXX^A486G^ displayed a significant overlap of genes involved in molecular pathways found to be associated to DAXX^A486G^ regulon-significant genes. The most prominent gene found to be over-expressed in mutant cells from sample GOE615, affecting the TP53 signalling at multiple levels, is the CDKN1A. Besides CDKN1A, the up-regulation of CCNG1, DDIT4, CAPG, TINF2, PPP1CB and DAXX result in enriched pathways for TP53 regulating transcription^32^. Moreover, the overexpression of CDKN1A is linked to TP53-dependent G1/S DNA damage checkpoint; TP53 regulating transcription of cell cycle genes; TP53-dependent G1 DNA damage response and TP53 regulating transcription of genes involved in G1 cell cycle arrest^33^. Notably, the DAXX gene itself was found to be over-expressed exclusively in sample GOE615, underlying the link from genetic variation to this specific progeria sample phenotype (Figure 6a).

Mutations in the DAXX gene have been found frequently in both telomerase-positive and alternative lengthening of telomeres (ALT) cells. Endogenous DAXX can localize to Cajal bodies, associated with the telomerase inducing DNA damage/telomere and ALT^34,35^.

In addition, when comparing the 27 significant regulon markers for MT-CYB^L236I^ with the genes over-expressed in mutant cells for the same mutation, it was evident that most of those genes participate in the same pathways; suggesting concordance in both analyses. Most of the induced genes are related to degradation of the extracellular matrix and specifically, transcriptional regulation by MECP2 via MET affecting diseases of signal transduction and sex-specific expression in autism and Rett syndrome^36^. Interestingly, five mitochondrial genes were found regulated by this mutation (MT-ATP6, MT-ATP8, MT-ND2, MT-ND4 and MT-CYB), resulting in enriched pathways, “mitochondrial respiratory chain complex I”, “mitochondrial proton-transporting ATP synthase complex” and “mitochondrial respirasome” (Figure 6b). As described above for DAXX gene, MT-CYB itself appears to be over-expressed only for sample GOE1360, providing a consistent evidence of linking variation heterogeneity to the phenotype for sample GOE1360. Mutations in MT-CYB have been reported to cause isolated complex III deficiency leading to a variety of symptoms including failure to thrive, exercise intolerance, cardiomyopathy and encephalomyopathy^37^. Some phenotypic variability in the gene has been reported in patients with additional complications of hearing loss, visual impairment, brain atrophy, cardiomyopathy and gastroparesis^38^.

## Discussion

In this study, we systematically compared the performance of two platforms for single-cell RNA-sequencing in primary dermal fibroblast cells by using a full-length approach demonstrating its utility in multi-omics analysis such as the ability of a genotype to show phenotypes diversity.

We combined two different instruments, the CellenONE for cell dispensing and the ICELL8 for fully automated RT and library preparation. This combination improved enormously the cell isolation quality step by sorting living or fixed cells based on cell sizing from nuclei or whole cells (3 to 500 µm in diameter), cell morphology and/or selecting cells by using one or more fluorescence labelling.

In our analysis, the performance of the novel platform allowed the investigation of more than 3,300 cells per run after executing quality control (QC) filtering on one ICELL8® 5,184 nano-well chip. Moreover, the combined platform displayed higher sensitivity in gene detection compared to the ICELL8 (Figure 2a and 2d). It was discovered that 30% of the genes detected in the novel platform were lncRNAs found across all patient samples investigated. Considering the whole human transcriptome, 2-3% of these nucleotide sequences can code for proteins, while the other 97% are ncRNAs that partially or completely lack the ability to be coded into proteins^39,40^. Because the chemistry used in this study is an oligo (dT) priming based cDNA synthesis and some of the lncRNAs are partially polyadenylated, many of these molecules can be detected on the single cell level using this approach. However, the coverage depth for the investigation of ncRNAs evaluating disease-markers or their effects on the transcriptomic context required a higher sequencing depth with at least 0.5 Mio reads/cell to improve the coverage depth.

The novel platform displays many of the markers found from the ICELL8 platform (nearly 97%) and it is also efficient in providing additional markers per sample by deeper coverage of the transcriptome (Figure 2d); thus, offering additional biological insight. Moreover, we showed that the chemistry applied covering transcripts in a full-length manner is essential for the investigation of transcriptional and genetic heterogeneity of one single cell.

The next challenge was connecting genetic regulation to phenotype in the six patient samples, and this was addressed by associating variants with gene expressions. However, variant detection for scRNA-seq data applying traditional methods do not work well, as they produce too many false positives. In order to minimize them and achieve a high specificity, we introduced filtering criteria based on (1) confirmation of variants with bulk RNA-seq data, and (2) discarding low depth variants. This improvement can be done when reads generated are covering whole transcripts with at least 300K reads per cell. Moreover, it is essential to corroborate variants detected on the scRNA-Seq with either bulk RNA-seq or DNA-seq approaches.

After applying the filtering criteria and sorting variants selectively expressed in each of the samples, between 94-756 variants were reported for each sample. From those variants detected, a regulon analysis was performed in order to evaluate connectivity of variations to marker-driven phenotypes by selecting markers with FDR<0.05. However, the performance of regulon analysis requires confirmation by differential expression analysis with mutant versus non-mutant cells. The definition of non-mutant cells is given by real wild-type cells or cells exhibiting low or no coverage at the variant position. Notably, the network of regulated genes found in the DEG analysis corroborates the markers proposed in the regulon; providing better knowledge and insights into molecular pathways of mutant cells driving phenotypes in the progeria samples.

It is important to note that these findings were done with only one variant for one specific gene in a specific sample and contribute partially to the phenotype diversity description. We reported two prominent pathways that were found to be altered for variation DAXX^A486G^ in GOE615, the TP53 regulating transcription and the DNA damage/telomere inducing alternative lengthening of telomeres via CDKN1A and DAXX. Regarding sample GOE1360, we reported variant MT-CYB^L236I^ contributing to regulation of several others mitochondrial genes (MT-ATP6, MT-ATP8, MT-ND2, MT-ND4 and MT-CYB) and regulation of MECP2 transcription via MET.

Here we proposed a new approach for scRNA-Seq that significantly improved the quality of selected single-cells based on excellent imaging features, allowing the exclusion of damaged cells generating artificial cell clusters, the analysis of whole cells independently of cell morphology or cell sizing as well as the use of nuclei. The chemistry used was a single-cell full-length RNA-seq that improves significantly the characterization of mutational and transcriptional heterogeneity. Finally, we demonstrated the applicability of this scRNA-Seq approach for determing accelerated ageing-associated pathway signatures and highlighting the possibility to identify additional variant-specific expression profiles that might influence the phenotypic presentation.

## Methods

### Study design

A total of eight primary dermal fibroblast cell cultures were used for the comparison of both cell-preparation approaches (ICELL8 library preparation and CellenONE cell extraction combined with ICELL8 library preparation).

### Cell culture of dermal Fibroblasts

Primary dermal fibroblast established from patient and respective controls were cultured in Dulbecco’s modified Eagle medium (DMEM, Gibco) supplemented with 10% fetal calf serum (FCS, Gibco), and antibiotics. Control samples GOE1303 and GOE1305 were purchased from Coriell Institute, New York, USA (GM02936C, GM00409D). GOE1309 were purchased from Coriell Institute, New York, USA (GM02548D) and GOE1360, GOE800, GOE486, GOE247 and GOE615 were provided by collaborators described in acknowledgements.

### Bulk RNA-sequencing

The start material (cell suspensions) used for single cell isolation were used for bulk sequencing (same source). Ca. 100,000 cells were used for RNA extraction using the Trizol Reagent (Thermo Fisher) according to manufacturer’s recommendations. RNA-seq libraries were performed using 100 ng total RNA of a non-stranded RNA Seq (TruSeq RNA Library Preparation Cat. N°RS-122-2001). Libraries were sequenced on the Illumina HiSeq 4000 (SE; 1 × 50 bp; 30-35 Mio reads/sample).

### Full-length single-cell RNA-seq using the ICELL8 system

The ICELL8 250v Chip was used with the full-length SMART-Seq ICELL8 Reagent Kit. Cell suspensions were fluorescent-labelled with live/dead stain, Hoechst 33342 and propidium iodide (NucBlue™ Cell Stain Reagent, Thermo Fisher Scientific) for 15 min prior to their dispensing into the a Takara ICELL8® 5,184 nano-well chip. CellSelect® Software (Takara Bio) was used to visualize and select wells containing single and live cells. Next, cDNA was synthesized via oligo-dT priming in a one-step RT-PCR reaction. P5 indexing primers for subsequent library preparation were dispensed into all wells receiving a different index, in addition to Terra polymerase and reaction buffer. Transposase enzyme and reaction buffer (Tn5 mixture) were dispensed to selected wells. P7 indexing primers were dispensed to wells. Final Illumina libraries were amplified and pooled as they were extracted from the chip. Pooled libraries were purified and size selected using Agencourt AMPure XP magnetic beads (Beckman Coulter) to obtain an average library size of 500 bp. A typical yield for a library comprised of ∼1300 cells was ∼15 nM. Libraries were sequenced on the HiSeq 4000 (Illumina) to obtain on average ∼ 0.3 Mio reads per cell (SE; 50bp).

### Full-length single-cell RNA-seq using the CellenONE-ICELL8 system

In the combined CellenONE-ICELL8 system, we used CellenONE to dispense single cells into the nano-wells of ICELL8 chip. The cell dispensing was carried out on a temperature-controlled target holder in a humidity-controlled chamber. Placement and rotational errors were adjusted using the built-in “Find Target Reference Point (FTRP)” function. The cell suspension (300 cells/µl) was used to first create a mapping that determine the cell size parameters based on cell types. For isolation of single human dermal fibroblasts, the following parameters were used: diameter 20-60 µm, elongation and circularity 1-1.5 µm excluding doublets and debris. After single cell dispensing, the nano-well chip was sealed and centrifuged. Next, the nano-well chip was placed in the iCELL8 cx system and proceeded with the RT and library preparation as described in the section above.

### Pre-processing of data from the ICELL8 and CellenONE-ICELL8 platform

Raw sequencing files (bcl) were converted into a single fastq file using Illumina bcl2fastq software (v2.20.0.422) for each platform. Each fastq file was demultiplexed and analyzed using the Cogent NGS analysis pipeline (CogentAP) from Takara Bio (v1.0). In brief, “cogent demux” wrapper function was used to allocate the reads to the cells based on the cell barcodes provided in the well-list files. Subsequently, “cogent analyze” wrapper function performed read trimming with cutadapt^41^(version 3.2), genome alignment to Homo sapiens genome GRCh38 using STAR^42^(version 2.7.2a), read counting for exonic, genomic and mitochondrial regions using featureCounts^43^(version 2.0.1), and utilizing Homo sapiens gene annotation version 103 from ENSEMBL and generating a gene matrix (https://www.ensembl.org/Homo_sapiens/Info/Index) with number of reads expressed for each cell in each gene. Raw gene matrices underwent quality control (QC) filtering for cells and genes using the following parameters: (a) for cells, only those with at least 10000 reads associated to at least 300 different genes, and (b) for genes, only those containing at least 100 reads mapped to them from at least 3 different cells.

### Gene marker detection

To determine gene markers expressed in specific samples, QC-filtered read count matrix was used as input to determine which genes were differentially expressed between cells from a particular sample and all other cells using Wilcoxon Rank Sum and Signed Rank Test, with p-values adjusted using Benjamini-Hochberg method. For the individual differential expression results, a rank score was calculated for each gene using the formula rank=-log_10_(p-value)*log_2_FC, where p-value is the raw p-value and log_2_FC is the log_2_ fold-change.

### Correlations of scRNA datasets and bulk RNA samples

Single-cell data sets were correlated for each sample individually for a select subset of genes. In brief, the markers in each sample from both scRNA-seq platforms were determined. The absolute rank scores from both platforms were summed for each sample individually, and the top 100 markers with highest absolute rank scores were used to correlate the ICELL8 and CellenONE-ICELL8 data sets using the Pearson correlation. Moreover, single-cell dataset pseudo-counts were calculated as the sum counts for all genes in cells from each particular sample, and the resulting pseudo-counts were combined with the bulk RNA-seq samples to a single gene matrix, normalized using scale sizes calculated by DESeq2^44^(version 1.32.0) and correlated for the same markers.

### Unsupervised cell cluster

Using the CogentDS R package, dimensionality reduction and visualization were performed with the t-SNE algorithm using the optimal principal components (PCs) chosen from elbow plots. In addition, unsupervised graph-based clustering was performed in order to cluster the cells into particular clusters. In brief, the indicated PCs were used to subset the normalized gene matrix, which was used to group the cells into clusters using the k-nearest neighbour algorithm (k=20) followed by a shared nearest neighbour algorithm and the “FindClusters” function from the Seurat R package using resolution=0.5 to associate the cells with the cluster^45^. For each graph-based clustering, gene markers were detected, and the top 30 genes were plotted in a heatmap.

### Variant calling in scRNA and bulk data

For the scRNA-seq data, aligned reads were separated based on their sample affiliation into 8 different alignment files, and variants were identified in those alignment files, as well as alignment files for bulk RNA samples, using the Genome Analysis Toolkit^46^(GATK v4.1.927). Using in-house R scripts utilizing the R package VariantAnnotation^47^(v1.38), the VCF files were merged together, and for each patient sample (GOE1360, GOE800, GOE486, GOE247, GOE615 and GOE1309), sample-specific scRNA-seq variants corroborated by bulk data and with a high variant and total depths were retrieved.

### Expression signatures of target variants

For each sample, a single sample-specific variation was selected, such that it had a maximum variant depth, as well as being nonsynonymous. For the target variations, the percentages of cells containing the mutation out of all cells in each sample were calculated, followed by a multiple regression analysis between average gene expressions per sample and the mutation percentages in each sample. The regression was calculated with the formula:

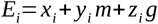

where *E*_*i*_ is the normalized average expression of each tested gene i, *z*_*i*_ *g* are the normalized average expressions of the gene affiliated with the target mutation per sample, and *y*_*i*_ *m* are the percentages of mutant cells, for which the p-value was calculated. From the regression analysis, p-values were calculated for all genes found to be markers in at least one sample, and corrected using the Benjamini-Hochberg. The normalized expressions of all QC-filtered cells were plotted in a heatmap for the top 30 genes with lowest adjusted p-values.

### Deregulated genes in mutant cells

using an in-house R script, the reads used to call each of the target variants were affiliated with their specific cell, such that cells with variation-containing reads were deemed “mutant cells” and others “non-mutant cells”. Similarly to the “Gene marker detection” method mentioned above, Wilcoxon Rank Sum and Signed Rank Test was used to find deregulated genes between mutant and non-mutant cells. The resulting genes were filtered for the initial value of absolute log_2_ fold-change >=0.25 (discarding small differences in expressions between cell groups) to allow a more lenient adjustment of the p-values using Benjamini-Hochberg method. Finally, the up- and downregulated genes were inserted into STRINGDB^48^ and REACTOME^49^ to uncover how the deregulated genes are connected to each other and what ontologies and/or pathways they belong to.

## Data availability

All sequencing data has been deposited in the Gene Expression Omnibus (GEO) at GSE179211. Any other relevant data are available from the authors upon reasonable request.

## Acknowledgements

We thank Michael Walter, Esra Kilic, Uwe Kornak, Barbara Leube, Thomas Schmitt-Melchelke for providing patient-derived fibroblasts. Wild-type (GM02936C and GM00409D) and BLM-deficient fibroblasts (GM02548D) were purchased from Coriell Institute, New York, USA. This work was supported by the Deutsche Forschungsgemeinschaft (DFG, German Research Foundation) under Research Group FOR 2800 “Chromosome Instability: Cross-talk of DNA replication stress and mitotic dysfunction”, SP5 and SPZ to B. Wollnik, Germany’s Excellence Strategy, Cluster of Excellence “Multiscale Bioimaging: from Molecular Machines to Networks of Excitable Cells” (MBExC; EXC 2067/1-390729940) to B. Wollnik, the SFB 1002 “Modulatorische Einheiten bei Herzinsuffizienz”, and the DZHK (German Center for Cardiovascular Research, Göttingen) to L. Cyganek and B. Wollnik.

## Author contributions

G. Salinas and B. Wollnik conceived the project and designed the experiments. K. Streckfuss Boemeke, L. Zelarayan and G. Salinas led the method development. F. Ludewig and S. Parbin led the experimental data production. O. Shomroni and M. Sitte led and performed the data analysis. J. Schmidt and G. Yigit contributed to experimental data and G. Salinas wrote the paper.

## Notes

https://www.ncbi.nlm.nih.gov/geo/query/acc.cgi?acc=GSE179211

### Competing Interest Statement

The authors have declared no competing interest.

